# MINI REVIEW: Statistical methods for detecting differentially methylated loci and regions

**DOI:** 10.1101/007120

**Authors:** Mark D Robinson, Abdullah Kahraman, Charity W Law, Helen Lindsay, Malgorzata Nowicka, Lukas M Weber, Xiaobei Zhou

**Affiliations:** Institute of Molecular Life Sciences, University of Zurich, CH-8057 Zurich, Switzerland; SIB Swiss Institute of Bioinformatics, University of Zurich, CH-8057 Zurich, Switzerland

## Abstract

DNA methylation, the reversible addition of methyl groups at CpG dinucleotides, represents an important regulatory layer associated with gene expression. Changed methylation status has been noted across diverse pathological states, including cancer. The rapid development and uptake of microarrays and large scale DNA sequencing has prompted an explosion of data analytic methods for processing and discovering changes in DNA methylation across varied data types. In this mini-review, we present a compact and accessible discussion of many of the salient challenges, such as experimental design, statistical methods for differential methylation detection, critical considerations such as cell type composition and the potential confounding that can arise from batch effects. From a statistical perspective, our main interests include the use of empirical Bayes or hierarchical models, which have proved immensely powerful in genomics, and the procedures by which false discovery control is achieved.

## 1 Introduction

Epigenomics can be defined as the genome-wide investigation of stably heritable phenotypes resulting from changes in a chromosome without alterations in the DNA sequence [1]. DNA methylation is the most well-studied epigenetic mark and notably, the enzymatic mechanism for mitotically copying methylation status is well understood [2], unlike the mechanism for maintaining chromatin state [3]. In this review, we focus on differential methylation (DM) for methyl groups added to cytosines in the CpG dinucleotide context, since this is the predominant form observed in differentiated mammalian cells [4]. However, some of the statistical methods and technologies discussed here can be applied more generally.

In the last decade, considerable progress has been made in (observationally) characterizing epigenetic phenomena across a wide spectrum of normal and disease states, predominantly due to large-scale explorative studies using emerging disruptive technologies, such as microarrays and large-scale sequencing of DNA [5, 6]. These studies have highlighted the important causal associations of DNA methylation with gene regulation and its potential in diagnosing or stratifying patients according to their combined genomic/epigenomic molecular state [e.g., 7]. Robust and efficient statistical and computational frameworks must be developed to facilitate interpretation of the growing masses of data. For CpG methylation, the main workhorse is treatment of DNA with sodium bisulphite [8], which preserves methylated cytosines while converting unmethylated cytosines to uracil. This transformation can allow high-throughput readouts, whether by microarray hybridization or DNA sequencing, to quantify the (relative) level of methylation.

While an individual’s (pathologically normal) genome is almost completely static in all cells, the epigenome is highly dynamic both in time (e.g., through development) and across cell types. Since the epigenome is a combinatorial assembly of regulatory factors (e.g., DNA methylation, histone modifications, non-coding RNAs, etc.), comprehensively profiling the epigenome is orders of magnitude more difficult than genome sequencing. Therefore, accurately measuring DNA methylation or other layers of the epigenome requires additional considerations to ensure that detected changes are not confounded with external factors, such as cell type.

Not surprisingly, the community has embraced consortium science to scale up data collection efforts. Prominent projects that involve large-scale profiling of DNA methylation include the ENCODE Roadmap Epigenomics Consortium [9], The Cancer Genome Atlas and International Cancer Genome Consortium [10, 11], the BLUEPRINT project [12] and the International Human Epigenome Consortium [13]. See Table 1 for further description.

**Table 1:**
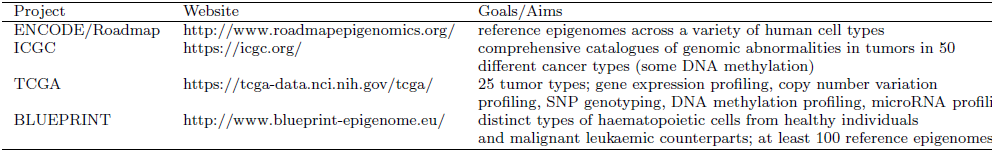
Active consortium science projects with DNA methylation data

## 2 Technologies

Present techniques for interrogating methylation fall into three categories: methylation-specific enzyme digestion, affinity enrichment and chemical treatment with bisulphite (BS). Techniques have been used in combination (e.g., enzyme digestion then BS, commonly known as RRBS; see 14), and with high-throughput readout. Early demonstrations were able to distinguish methylcytosine from cytosine with third-generation technologies [15], but no commercially viable offering has yet appeared. Methylation profiling techniques vary in resolution from low (*∼*100-200 base pair) to high (individual CpG sites) and their costs vary widely. Each platform has its own limitations related to cost, resolution, scalability and the amount of starting DNA required [16, 14, 17]. For example, enzyme digestion studies remain dependent on the location and frequency of enzyme restriction sites; the prominent BS-based microarray platform is only available for human; the sensitivity of enrichment approaches depends on CpG density, while genome-scale sequencing-based BS methods are costly and require considerable computing resources [18]. Depending on the biological question and resources available, a platform may be selected based on these tradeoffs.

Notably, BS-based methods cannot distinguish between methylcytosine and other variants, such as hydroxymethylcytosine [19], although additional treatment steps can readily allow this [20]. The methods we discuss below are agnostic to this technicality, aside from specific biological questions regarding the interplay between methylation states. Another biological phenomenon we sidestep in this review is methylation in non-CpG contexts, shown to be prominent in pluripotent cells [4]. Interestingly, a recent report using whole genome BS-seq across various cell types found that CpG methylation is only “dynamic” in approximately 20% of sites [6], suggesting that BS-seq could be more directed. Enzyme digestion (and size selection) with BS-seq is already a favored method to reduce sequencing depth, but is difficult to tailor to specific genomic regions. An alternative reduced complexity strategy is to first capture fragments of interest, for example by using the Agilent SureSelect system [21].

A popular, cost-efficient and scalable technology for profiling DNA methylation on a “genome-scale” is the Illumina 450k microarray. The platform can be thought of as genotyping BS-treated DNA to reveal the relative proportion of methylated and unmethylated alleles [22]. For every CpG site, measurements are either made with two separate physical beads (Type I) or through a single bead across two fluorescence channels (Type II); properties of these probe types are vastly different and require careful normalization [23, 24].

With the decreasing costs of single-base resolution DNA methylation data, enrichment techniques that capture methylated DNA fragments appear to have gone somewhat out of favor. Methylated DNA immunoprecipitation (MeDIP) or methyl-binding domain enrichments share many features of chromatin immunoprecipitation experiments. However, they are plagued by enrichment biases caused associated with CpG density, some of which can be fixed *in silico* [18, 25, 26]. Recently, a combination approach of MeDIP-seq and methylation-sensitive restriction enzyme sequencing (MRE-seq) has become available, promising to quickly compare methylomes at lower cost [27].

Table 2 summarizes the methods reviewed and gives brief details on some of the important features: i) data type; ii) ability to define regions; iii) support for covariate adjustment; iv) statistical tests used.

**Table 2:**
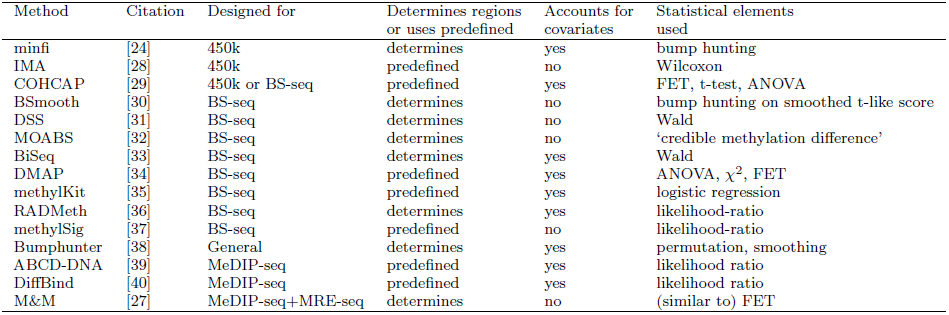
Recent methods for differential methylation detection.

## 3 Experimental design

Ultimately, the same experimental design concepts that apply broadly to any scientific investigation, such as sampling, randomization and blocking, are assumed. Excepting single-cell DNA methylation studies [e.g., 41], it is crucial to remember that every experimental unit represents a *population* of cells. This implies that a consensus methylation estimate of 50% could mean 50% of the alleles are methylated in all cells (e.g., allele-specific methylation) or 50% of the cells are fully methylated (e.g., mixtures of cell types), or any combination thereof. Only BS-seq data can properly decompose this information, and at the same time infer allele-specific patterns, using the methylation status from individual DNA fragments [42, 43, 44]. But, there are limits: since small fragments are observed, it remains challenging to relate the allelic methylation status at one loci to another genomically distant loci without additional haplotype information [45].

In contrast to genome sequencing studies, collecting relevant populations of cell types is important for epigenome profiling projects. Many population-scale profiling studies may consider using readily accessible bodily fluids such as blood, which represents a rich milieu of cell types that may vary in composition across the experimental units being studied. If the cell types and cell surface markers of interest are known, it may be beneficial to first sort cells into subpopulations and profile each individually [46]. Doing so will give a more focused interrogation of methylation and improved signal over noise. However, there are many situations where pre-sorting is not possible. Importantly, profiling mixtures of cell types and looking for changes in DNA methylation can be misleading when the cell composition is associated with an external factor, such as age of the patient [47]. However, there are now various emerging computational techniques to deconvolute the cell composition signals *in silico* (see Section 6).

Another design consideration for BS-seq experiments is whether money is better allocated towards deeper sequencing or additional replicates. Because of the local smoothing frameworks available for methylation measurements (e.g., BSmooth, 48), it is considered better to sequence additional replicates than to gather deep information on fewer samples.

## 4 Finding differential loci

We first focus on the methodology for discovering individual differentially methylated CpG **sites** for single-base resolution assays. BS-seq data can be summarized as counts of methylated and unmethylated reads at any given site. Many early BS-seq studies profiled cells without collecting replicates and used Fisher’s exact test (FET) to discern DM [49]. While this strategy may be sufficient for comparing cell lines, we stress that the use of FET should be generally avoided; most systems have inherent biological variation and FET does not account for it. For example, in a two-condition comparison, FET requires the data to be condensed to counts for each condition, completely ignoring the within-condition variability. This will underestimate variability and overstate differences, leading to a high false positive rate. Likewise, using the binomial distribution, e.g. within a logistic regression framework (e.g., methylKit; 35), does not facilitate estimation of biological variability unless an overdispersion term is used. While BSmooth uses a “signal-to-noise” statistic to quantify DM evidence at individual CpG sites, it is not used directly for inference of differential sites (more details in Section 5).

The most natural statistical model for replicated BS-seq DNA methylation measurements is beta-binomial. Conditional on the methylation proportion at a particular site, the observations are binomial distributed, while the methylation proportion itself can vary across experimental units (e.g., patients), according to a beta distribution. It is therefore no surprise that beta-binomial assumptions are made in several recent packages, such as BiSeq [33], MOABS [32], DSS [31], RAD-Meth [36] and methylSig [37]. Similarly, empirical Bayes (EB) methods fit naturally for modeling and inference across many types of genomic data, including DNA methylation assays. MOABS and DSS both implement hierarchical models and use the full dataset to estimate the hyperparameters of the beta distribution; RADMeth, BiSeq and methylSig use standard maximum likelihood without any moderation. While BiSeq and RADMeth do not moderate parameter estimates, they provide facilities for complex designs through design matrices, which MOABS, DSS and methylSig do not currently offer. Inference for parameters of interest (i.e., changes in methylation) are conducted using standard techniques, such as Wald tests (DSS, BiSeq) and likelihood ratio tests (RADMeth, methylSig). Notably, MOABS introduces a new metric, called credible methylation difference, which is a conservative estimate of the true methylation difference, calibrated by the statistical evidence available.

DNA methylation arrays, such as Illumina’s 27k or 450k array, give fluorescence intensities that quantify relative abundance of methylated and unmethylated loci, in contrast to the count-based modeling assumptions for BS-seq based profiling. In particular, the data used for downstream analyses can be either i) log-ratios of methylated to unmethylated intensities, or ii) the beta-value, which gives the ratio of the methylated to the total of methylated and unmethylated intensities. Previous comparisons suggest that statistical inferences based on log-ratios are preferred [50], perhaps not surprisingly since they can rely on earlier successful moderated statistical testing strategies (e.g., limma; 51). Much of the recent effort for the 450k array has been dedicated to normalization and filtering (e.g., 24, 52) and various options for inferring DM sites from 27k/450k array data have been proposed. To test for DM, IMA proposes Wilcoxon rank-sum tests on beta-values [28]. COHCAP operates either on methylation array data or BS-seq data, using beta-values or methylation proportions as input; they offer FET (see comment above), t-tests and ANOVA analyses (without moderation), depending on the study design [29]. Ultimately, we speculate that moderated t/F-statistics on the normalized log-ratios of intensities should perform well.

## 5 Finding differential regions

Although there are occasions when researchers are interested in relating single CpG sites to a phenotype [e.g., 53], often differentially methylated regions (DMRs) are a more predictive feature. Another advantage is that while differences at any individual site may be small, if they are persistent across a region, statistical power to detect them may be greater. Methods that *operate* on predefined regions must be distinguished from those that *define* regions of DM. The latter is considerably more difficult because ensuring control of the false discovery rate (FDR) at the region-level is non-trivial; in particular, FDR control at the site-level does not give a direct way to region-level control when the region itself is also to be defined [54].

Therefore, the most straightforward approach is to use predefined regions, such as CpG islands, CpG shores, UTRs and so on; statistical testing can be conducted fairly routinely at a region-level. Many of the packages mentioned above, such as IMA, COHCAP, DMAP, methylSig and methylKit, do exactly this. A special case is DMAP, which can operate on fragments (using the sampled MspI-digested fragments as the region of interest) or according to predefined regions [34].

There are now many approaches for *defining* DMRs. For example, Bumphunter can be applied quite generally across data types [38], perhaps after transformation in the case of count data. Notably, it also integrates a surrogate variable analysis [55] to simultaneously account for potential batch effects while permutation tests are used to assign FDR at the region-level; users should set a smoothing window size and a threshold on the percentile of the smoothed effect sizes (or t-statistics) [38]. Similarly, BSmooth searches for runs of smoothed absolute t-like scores beyond a threshold, however, does not suggest a permutation strategy to control region-level FDR. From the same authors, minfi wraps bumphunting into the suite of methods available for Illumina 450k arrays; in addition, they provide a module for *block finding*, which is essentially bumphunting with a much greater window size (e.g., 250kbp) [24]. BiSeq proposes, via a Wald test statistic from the beta-binomial regression fit, a hierarchical testing strategy that first considers target regions and controls error using a cluster-wise weighted FDR strategy [33]; secondly, the differential clusters are trimmed using a second stage of testing, analogous to methods used for spatial signals [56]. A clustering method, A-clust, proposes first to cluster CpG sites according to correlation in methylation signal across samples; within the clusters, associations can be modeled with correlated error and fit using a generalized estimating equation framework [57]. The DSS authors simply set some thresholds on the P-values, number of CpG sites and length of regions, but they do not pursue FDR control [31]. The MOABS authors suggest grouping DM sites into DMRs using a hidden Markov model or alternatively testing of predefined regions, but no specific details are given. RADMeth proposes a transformation of P-values (from a likelihood ratio test) into a weighted Z test that builds in the correlation of neighboring probes [58, 36]; the same adjustment, also known as the Stouffer-Liptak test, is used in the eDMR tool [59]. Also used in the context of DMR detection for combining spatially correlated P-values, but applicable more generally, is a tool called comb-p [60].

Enrichment assays, such as MeDIP-seq, are by their very nature of capturing fragments, only capable of finding *regions* of DM. Packages for considering enrichment data, including MEDIPS [61, 62], ABCD-DNA [39] and DiffBind [40], compare relative abundance of fragment counts by repackaging RNA sequencing statistical frameworks. In a related assay, the M&M algorithm models normalized bin-wise methylated counts (MeDIP-seq) and unmethylated counts (MRE-seq) as jointly Poisson distributed with a shared parameter [27]. Analogous to the FET, the Poisson model does not account for biological variability [27].

## 6 Reducing the impact of batch, cell type composition or other confounding effects

Researchers need to carefully design studies that associate phenotypes with DNA methylation. Some aspects, such as cell type composition, cannot be readily controlled by design; patients and therefore individual DNA samples simply differ in their cell type composition. A recent report has highlighted that many of the DNA methylation markers that have been associated with age are actually driven by age-related changes in cell composition [47]. Whole blood is a mixture of several cell types; using an independent dataset of methylation profiles of the dominant cell types (Monocytes, CD4+ and CD8+ T cells, Granulocytes, B cells, natural killer cells) from flow sorting, patient profiles were deconvoluted using a reimplementation of the Houseman algorithm [47, 46]. From the methylation profiles of pure cell populations, cell-type-specific markers are selected and then used to “calibrate” a regression model that associates methylation observations to a response of interest [46, 63]. Of course, this approach requires advance knowledge of the dominant cell types and methylation profiles for them, preferably across multiple replicates to seed the deconvolution algorithm with appropriate methylation markers. However, a recent study has highlighted that advanced statistical modeling can correct for cell type composition without the need for pure-cell profiles; starting from uncorrected standard model fits, the method regresses principal components within a linear mixed model until control for the inflation of the test statistics (e.g., relative to a uniform distribution of P-values) is achieved [64].

## 7 Discussion

In this review, we briefly explored the various methodologies available for deciphering DMRs across the main data types and highlighted some of the common themes and current challenges. The tradeoffs made by method developers are apparent. In fact, it’s a lot to ask of a single statistical framework to do everything: moderate parameter estimates using genome-wide information or accurately and robustly smooth local estimates, accommodate low coverage data, account for batch effects and cell type composition, allow complex experimental designs and accurately control FDR at the site- and/or region-level. In addition, identification of DMRs is only the discovery step; validating these detections, perhaps by associating them with other biological outcomes *in silico* requires additional frameworks, some of which have already been integrated alongside the packages reviewed here.

On the statistical and computational side, the field is moving fast and several advanced methods have been proposed. One of the next challenges will be to comprehensively compare method performance, in terms of statistical power, ability to control FDR, robustness and scalability to large datasets and large studies. Representative simulation frameworks will be fundamental for this task. Given the large number of methods available, this will already be a large undertaking. To avoid bias, these comparisons should be done either independently of the method development process, or collectively with all method developers. Advanced deconvolution algorithms and batch effect removal strategies are, at present, targeted at 450k array data. The development and vetting of similar techniques that can be readily applied to count data, such as BS-seq data, are well underway [65, 66].

## Author Contributions

MDR drafted the text with contributions from all co-authors: AK, CWL, HL, MN, LMW and XZ. All authors read and approved the final manuscript.

## Acknowledgement

We would like to thank followers on Twitter (@timtriche, @PeteHaitch, @brent p) and especially Peter Hickey for suggesting additional citations and/or carefully reading an earlier draft.

